# Integrated Metabolomic, Proteomic, and Phosphoproteomic Profiling Reveals CAMKK2-Dependent Regulation of Cell Cycle and Nucleotide Metabolism in Gastric Cancer

**DOI:** 10.64898/2026.04.23.720293

**Authors:** Mohd Altaf Najar, Nikita Choudhary, Nidhi Dwived, Prashant Kumar Modi

## Abstract

Gastric cancer is driven by aberrant kinase signaling that supports uncontrolled proliferation and metabolic adaptation. Calcium/calmodulin dependent protein kinase kinase 2 (CAMKK2) is overexpressed in gastric cancer; however, its role in regulating metabolic programs that sustain tumor growth remains incompletely understood. In this study, we employed an integrated multi-omics approach with a primary focus on untargeted metabolomics to investigate the consequences of CAMKK2 inhibition in gastric cancer cells. Pharmacological inhibition of CAMKK2 using STO-609 in AGS cells resulted in significant suppression of proliferation, clonogenic growth, migration, and invasion, accompanied by pronounced nuclear abnormalities and multinucleation indicative of mitotic defects. Global metabolomic profiling revealed extensive and time-dependent metabolic reprogramming following CAMKK2 inhibition, characterized by a marked depletion of nucleotide intermediates, including purine and pyrimidine metabolites required for DNA synthesis. Pathway enrichment analysis highlighted suppression of nucleotide metabolism, lipid metabolism, and central carbon metabolic pathways, indicating a broad impairment of biosynthetic capacity. Integration with proteomic and phosphoproteomic datasets demonstrated that metabolic alterations were accompanied by downregulation of DNA replication machinery and attenuation of kinase signaling pathways governing cell cycle progression. Protein metabolite interaction and docking analyses further supported functional coupling between nucleotide metabolites and key replication-associated enzymes, revealing disruption of metabolite enzyme interactions upon CAMKK2 inhibition. Collectively, these findings identify CAMKK2 as a critical regulator of metabolic programs that support DNA replication and cell cycle progression. Its inhibition induces replication stress through coordinated depletion of nucleotide pools and disruption of replication-associated signaling, leading to impaired proliferation and mitotic failure. These results highlight CAMKK2 as a potential therapeutic target for exploiting metabolic vulnerabilities in gastric cancer.

## Introduction

Gastric cancer (GC) remains one of the most lethal malignancies worldwide, ranking as the fifth most frequently diagnosed cancer and the fourth leading cause of cancer-related mortality globally [1–3]. Its incidence shows marked geographic variation, strongly influenced by environmental and lifestyle factors, including dietary habits and Helicobacter pylori infection [4, 5]. Despite improvements in early detection and therapeutic strategies, the prognosis of advanced gastric cancer remains poor, largely due to an incomplete understanding of the molecular mechanisms that drive tumor progression and therapeutic resistance [6, 7]. At the molecular level, gastric cancer is characterized by widespread dysregulation of oncogenic signaling pathways, including aberrant activation of receptor tyrosine kinases and downstream MAPK and PI3K/AKT pathways [8–10]. A defining feature of gastric tumorigenesis is the disruption of cell cycle control, particularly at the G1-S transition, where alterations in Cyclin D1, RB1, and E2F-regulated transcriptional programs promote uncontrolled proliferation [11–13]. These processes are tightly regulated by kinase signaling networks that integrate extracellular cues with intracellular proliferative and metabolic programs.

Protein kinases play central roles in regulating cellular proliferation, survival, and metabolism, and their dysregulation is a hallmark of cancer [14, 15]. In gastric cancer, aberrant activation of kinases such as EGFR, HER2, KRAS, RAF1, and MAPKs has been extensively documented [10, 16]. In addition to these canonical pathways, emerging evidence highlights the role of calcium/calmodulin-dependent kinases in cancer biology [17, 18]. Among these, calcium/calmodulin-dependent protein kinase kinase 2 (CAMKK2) has gained attention as a key regulator of both signaling and metabolic processes [19–21].

CAMKK2 is a serine/threonine kinase that acts as an upstream regulator of several signaling cascades through activation of downstream kinases, including CAMKI, CAMKIV, and AMP-activated protein kinase (AMPK) [22, 23]. Through these pathways, CAMKK2 has been implicated in the regulation of transcription, energy homeostasis, and cell cycle progression [24, 25]. Importantly, CAMKK2 has been shown to be overexpressed in multiple cancer types, including gastric, prostate, and liver cancers, where it contributes to tumor growth and metastasis [20, 26]. Notably, CAMKK2 signaling exhibits strong context dependency across cancer types [27]. In prostate cancer, CAMKK2 promotes tumor growth through AMPK-mediated metabolic reprogramming and increased glucose uptake,[28] whereas in ovarian cancer, it has been shown to activate AKT-dependent survival pathways [29, 30]. These findings suggest that CAMKK2 functions as a critical integrator of signaling and metabolic networks, enabling cancer cells to adapt to proliferative and energetic demands.

Recent advances in cancer biology have highlighted the importance of metabolic reprogramming as a hallmark of tumor progression [31, 32]. In particular, nucleotide metabolism plays a central role in supporting rapid cell proliferation by providing essential building blocks for DNA replication [33, 34]. Cancer cells exhibit increased reliance on de novo purine and pyrimidine biosynthesis pathways to sustain nucleotide pools required for S-phase progression [35–37]. Perturbations in nucleotide metabolism can lead to replication stress, characterized by stalled replication forks, genomic instability, and defective cell division [38–40].

Importantly, emerging evidence suggests that metabolic pathways are tightly coupled with signaling networks to coordinate DNA replication and cell cycle progression [41, 42]. However, the molecular mechanisms linking kinase signaling to nucleotide metabolism in gastric cancer remain poorly defined. While previous studies, including our own, have demonstrated that CAMKK2 promotes gastric cancer cell proliferation through MAPK–CDK–MCM signaling [41, 42], a comprehensive understanding of how CAMKK2 integrates phosphorylation-dependent signaling with metabolic pathways that sustain DNA replication has not been established. In particular, it remains unclear whether CAMKK2 directly regulates nucleotide metabolism and how disruption of CAMKK2 signaling impacts the coupling between metabolic availability and replication machinery. Given the central role of nucleotide metabolism in maintaining genomic integrity and proliferative capacity, elucidating this connection is critical for understanding CAMKK2-driven oncogenic processes.

To address this gap, we employed an integrated multi-omics approach combining untargeted metabolomics, quantitative proteomics, phosphoproteomics, and functional assays to systematically characterize CAMKK2-dependent regulatory networks in gastric cancer cells. Using pharmacological inhibition of CAMKK2 in AGS cells, we sought to define how CAMKK2 coordinates kinase signaling and metabolic pathways to sustain DNA replication and cell cycle progression, and to elucidate the mechanisms by which its inhibition induces replication stress and proliferative defects.

## Materials and Methods

### Reagents and Chemicals

The human gastric cancer cell lines AGS were obtained from American Type Culture Collection (ATCC, Manassas, VA), USA. Dulbecco‘s Modified Eagle Medium-high glucose ((DMEM, Cat#12100046)), fetal bovine serum (FBS, Cat#10270-106), and 100X antibiotic/antimycotic solution (Cat#15240-062), Trypsin EDTA (Cat#25200-056), were purchased from Gibco, Thermo Fischer Scientific, USA, and phosphate-buffered saline (PBS, Cat#TL101), from HiMedia, India. All plasticware for cell culture was procured from Nunc.STO-609 (7-Oxo-7H-benzimidazo[2,1-a]benz[de]isoquinoline-3-carboxylic acid acetic acid), a CAMKK2 inhibitor, was purchased from Santa Cruz Biotechnology (Cat# sc-202820), USA. BisBenzimide H33342 (HOECHST, Cat#B2261), propidium iodide (PI, Cat#P4170), LC–MS grade methanol (Cat# 34860), acetonitrile (Cat# 34967), water (Cat# W4502), and formic acid (Cat# 56302) were procured from Sigma-Aldrich,Pierce™ BCA protein estimation assay kit (Cat# 23225).

### Cell culture

AGS gastric cancer cells were obtained from the American Type Culture Collection (ATCC, Manassas, VA, USA). Cells were maintained in a humidified incubator at 37 °C with 5% CO and cultured in Dulbecco’s Modified Eagle Medium (DMEM) supplemented with 10% fetal bovine serum (FBS) and 1× antibiotic antimycotic solution.

### Cell Culture and Treatments

Cell culture experiments were carried out with AGS gastric adenocarcinoma cells (ATCC® CRL-1739™), procured from the National Centre for Cell Sciences (NCCS), Pune, India. Cells were maintained in Dulbecco’s Modified Eagle Medium (DMEM-high glucose) supplemented with 10% fetal bovine serum (FBS) and 1% antibiotic/antimycotic solution at 37°C in a humidified atmosphere containing 5% CO. For experimental treatments, cells were seeded at a density of 1 × 10 cells per 10-cm dish or 3 × 10 cells per well in 6-well plates and allowed to adhere overnight. The CAMKK2 inhibitor STO-609 was dissolved in dimethyl sulfoxide (DMSO) to prepare a stock solution and further diluted in complete culture medium to achieve the desired working concentration. A final concentration of 0.1% DMSO was maintained in all experimental groups, including vehicle controls. Cells were treated with the CAMKK2 inhibitor at a final concentration of 18.5μM. Vehicle-treated cells (0.1% DMSO) served as controls. At the end of treatment, cells were washed with ice-cold phosphate-buffered saline (PBS) and processed immediately for metabolite extraction.

### Metabolite Extraction

Intracellular metabolites were extracted using a modified biphasic solvent system composed of acetonitrile, methanol, and water (2:2:1, v/v/v). Following treatment with the CAMKK2 inhibitor for 48 hr, AGS cells were rapidly washed twice with ice-cold 1× PBS to eliminate residual media components. Cells were then scraped in chilled Milli-Q water and immediately snap-frozen in liquid nitrogen for 10 min to halt enzymatic activity and preserve metabolite integrity. Cell disruption was performed on ice using a probe sonicator (Q-Sonica, Cole-Parmer) to ensure efficient lysis while minimizing thermal degradation. Total protein concentration in each lysate was quantified to allow normalization across samples prior to metabolite extraction. Protein estimation was carried out using a bicinchoninic acid (BCA) assay kit (Thermo Fisher Scientific, USA) and verified by SDS-PAGE followed by Coomassie Brilliant Blue staining to confirm equal loading. For extraction, aliquots corresponding to 100 μg of total protein were mixed with the pre-chilled extraction solvent. Samples were briefly incubated at room temperature, vortexed vigorously, and subjected to sonication in an ultrasonic water bath to enhance metabolite release. The mixtures were centrifuged at 12,000 × g for 15 min at 4°C to separate insoluble material. The resulting supernatant containing soluble metabolites was carefully transferred to fresh tubes. Extracted metabolites were evaporated to dryness using a SpeedVac concentrator (Thermo Fisher Scientific, USA) and stored at 20°C until subjected to downstream mass spectrometry based metabolomics profiling.

### LC-MS/MS-Based Untargeted Metabolomics

Untargeted metabolomic analysis was performed using a Q-TRAP 6500 hybrid triple quadrupole/linear ion trap mass spectrometer (AB SCIEX, USA) coupled to an Agilent Infinity II 1290 ultra-high-performance liquid chromatography (UHPLC) system (Agilent Technologies, USA). Instrument operation and data acquisition were managed using Analyst software (version 1.6).Metabolite separation was carried out on a reverse-phase C18 column (Zorbax Eclipse RRHD, 2.1 × 150 mm, 1.8 μm particle size; Agilent Technologies, USA). The chromatographic mobile phases consisted of solvent A (0.1% formic acid in Milli-Q water) and solvent B (0.1% formic acid in 90% acetonitrile). Dried metabolite extracts were reconstituted in solvent A and supplemented with epicatechin as an internal standard (final concentration: 862 nM). A volume of 10 μL was injected for each analysis.The gradient elution program was optimized as follows: 2% solvent B from 0-1 min; increased to 30% B at 10 min; raised to 60% B at 11 min; ramped to 95% B between 13-17 min to facilitate elution of non-polar metabolites; and finally returned to 2% B from 17.2–20 min to allow column re-equilibration.

Mass spectrometric acquisition was conducted using an enhanced mass spectra–information dependent acquisition-enhanced product ion (EMS-IDA-EPI) workflow. Full-scan MS data (EMS mode) were collected in low-mass mode over an m/z range of 50–1,000 Da. The scan cycle was segmented into defined mass windows (50–102.87 Da with 0.0053 s dwell time; 102.87–308.63 Da with 0.0206 s; and 308.63–1,000 Da with 0.0691 s) to ensure comprehensive coverage. The scanning rate was maintained at 10,000 Da/s with a dynamic fill time of 250 ms and a linear ion trap (LIT) fill time of 10 ms. Dynamic background subtraction was enabled to enhance signal specificity.

During IDA, the five most intense precursor ions detected in EMS (MS1) mode were automatically selected for fragmentation. MS/MS spectra were acquired in EPI mode using high-energy collision activated dissociation (CAD). Data acquisition was performed separately in positive and negative electrospray ionization modes with ion spray voltages of +4,500 V and - 4,500 V, respectively. The source temperature was maintained at 450°C. Compound-dependent parameters included a declustering potential (DP) of +75 V (positive mode) and −75 V (negative mode), and a collision energy (CE) of +40 V and −40 V, respectively, with a collision energy spread of ±25 V applied in both polarities.For metabolomic profiling, three independent biological replicates were analyzed, with three technical replicates per biological replicate. Blank solvent injections were introduced after every triplicate set to monitor system performance and minimize carryover.

### Processing of Untargeted Metabolomics Data

Raw mass spectrometry data files (.wiff format) were converted into .mzML format using MSConvert from the ProteoWizard suite [43]. The converted files were subsequently processed using MZmine version 2.53 [44]. The workflow and processing parameters applied in MZmine are described below. Peak detection at both MS1 and MS/MS levels was performed using a centroid-based algorithm. The minimum intensity thresholds were set to 1.0E3 for MS1 and 1.0E1 for MS/MS signals. MS/MS feature extraction was carried out using the peak builder module with an m/z tolerance window of 0.5 Da and a retention time window of 2 min. The detected centroid MS/MS peaks were further refined using the peak extender module with a minimum peak intensity threshold of 1000 (1.0E3). Chromatographic peak deconvolution was conducted using the noise amplitude algorithm, applying a noise threshold of 1.5E2 and a minimum peak height of 1.0E3. MS/MS pairing during deconvolution was performed with a mass tolerance of 0.1 Da and a retention time tolerance of 1 min. Isotopic peak grouping was executed using the isotope peak grouper function with a maximum charge state of 4, an m/z tolerance of 0.25 Da, and a retention time tolerance of 0.2 min. Feature alignment across biological and technical replicates was performed using the Join Aligner module, employing a mass tolerance of 0.05 Da and a retention time tolerance of 0.5 min. Alignment weighting parameters were set at 75% for m/z and 25% for retention time. To account for missing values, gap filling was applied using a mass tolerance of 0.05 Da and a retention time window of 0.5 min. Duplicate feature removal was subsequently carried out using the duplicate peak filter option, considering features with identical charge states and applying an m/z tolerance of 0.05 Da and a retention time tolerance of 0.2 min. The finalized peak list was exported in GNPS-FBMN format which resulted in .quant (CSV) containing MS1 and MS2 data and .mgf containing MS2 data information along with corresponding peak intensity and area values. The features file (.mgf) was further subjected to Systematical Error Removal using Random Forest (SERRF) analysis using R- script for quantile normalization. Principal component analysis (PCA) was conducted to assess clustering patterns between biological samples and blanks, thereby confirming analytical reproducibility and overall run quality (Ashokan et al., 2021).

### Database Search and Metabolite Annotation

Metabolite identification was performed using the MS2Compound tool (https://sourceforge.net/projects/ms2compound/) [45], enabling annotation at both precursor (MS1) and fragment (MS/MS) levels. An in-house reference library was employed as the primary search background, constructed using metabolite entries curated from the Human Metabolome Database (HMDB) [46]. For fragment-level annotation, theoretical fragmentation spectra generated using Competitive Fragmentation Modeling for Metabolite Identification (CFM-ID) were used as the reference database. Precursor-based searches incorporated m/z values, charge states, and unique feature identifiers as input parameters. MS/MS-based searches were conducted using the “.mgf” files generated from both positive and negative ionization modes. A precursor mass tolerance of 0.05 Da was applied consistently for both MS1 and MS/MS level searches. For MS/MS matching, a fragment mass tolerance of 0.5 Da was implemented under medium-energy collision conditions, and metabolite assignments required a minimum of two fragment ion matches to be considered valid. Candidate metabolites were ranked based on their computed mS-Score, and the highest-scoring match was assigned to each detected m/z feature for downstream analysis.

### Statistical Analysis

Processed metabolomics data generated from MZmine 2.53 were subjected to downstream statistical analysis using MetaboAnalyst6.0 using RaMP-DB pathway overrepresentation analysis [47]. Fold-change (FC) calculations and multi-group statistical testing of the MS/MS-aligned peak list generated from MZmine. For inferential statistics, a one-way analysis of variance (ANOVA) was applied across experimental groups. To correct for multiple hypothesis testing, p-values were adjusted using a false discovery rate (FDR). Metabolites meeting the predefined thresholds for fold change and FDR-adjusted significance were considered differentially regulated and were subjected to downstream pathway and enrichment analyses.Furthermore, integrated metabolite protein pathway analysis was conducted by combining metabolomics data with differentially regulated proteomic datasets generated previously enabling joint pathway interpretation at both metabolic and proteomic levels.

### LC–MS/MS–Based Quantitative Proteomics

Cells were harvested at ∼70% confluency, washed with ice-cold PBS, and lysed in TEABC buffer containing 2% SDS supplemented with phosphatase inhibitors (sodium orthovanadate, sodium pyrophosphate, and β-glycerophosphate). Protein concentration was determined using a BCA assay, and 800 μg of total protein from each sample was subjected to reduction (DTT) and alkylation (IAA). Proteins were precipitated using chilled acetone to remove SDS and subsequently digested overnight with sequencing-grade modified trypsin. The resulting peptides were labeled using TMT 10 plex reagents, with control samples assigned to reporter ions 126, 127N, 128N, and CAMKK2 inhibitor treated samples labeled with 129N, 130N, 130C. After confirming labeling efficiency, equal amounts of peptides were pooled and subjected to basic pH reverse-phase fractionation. Peptides were separated on an XBridge C18 column using a linear gradient (3–50% acetonitrile in TEABC, pH 8.5), and 96 fractions were concatenated into 12 pooled fractions prior to LC–MS/MS analysis. NanoLC-MS/MS analysis was performed on an Orbitrap Fusion Tribrid mass spectrometer coupled to an EASY-nLC 1200 system. Peptides were first loaded onto a C18 trap column and then separated on a 50 cm C18 analytical column at a flow rate of 300 nL/min using a 90 min gradient of 5-35% solvent B (80% acetonitrile in 0.1% formic acid). MS data were acquired in data-dependent acquisition mode. Full MS scans were collected in the Orbitrap (resolution 120,000 at m/z 200; scan range 400–1600 m/z), followed by HCD fragmentation of the most intense precursor ions (charge states 2–6) using a top-speed method (3 s cycle time). MS/MS spectra were acquired at a resolution of 30,000 with dynamic exclusion set to 30 s.

Raw files were processed using Proteome Discoverer (v2.2) and searched against the human RefSeq database using SEQUEST and Mascot algorithms. Carbamidomethylation of cysteine and TMT labeling at peptide N-termini and lysine residues were set as fixed modifications, while methionine oxidation and protein N-terminal acetylation were specified as variable modifications. Mass tolerances were set to 10 ppm for precursor ions and 0.05 Da for fragment ions, allowing one missed cleavage. Peptide-spectrum matches were filtered to a false discovery rate (FDR) of 1%.

### Quantitative Phosphoproteomics Analysis

AGS cells were cultured to ∼70% confluence and treated with the CAMKK2 inhibitor STO-609 or vehicle control for 2 h prior to harvesting. Cells were washed with ice-cold PBS and lysed in urea-based lysis buffer (8 M urea, 75 mM NaCl, 50 mM Tris-HCl pH 8.0, 1 mM EDTA) supplemented with phosphatase inhibitors. Lysates were sonicated on ice, clarified by centrifugation, and protein concentration was determined using a BCA assay. Equal amounts of protein (1 mg per sample) were reduced with dithiothreitol, alkylated with iodoacetamide, and diluted with TEABC to reduce urea concentration prior to overnight digestion with TPCK-treated trypsin (1:20 enzyme-to-protein ratio) at 37°C. Peptides were desalted using C18 cartridges and dried prior to labeling. For quantitative analysis, peptides from control and STO-609-treated samples (biological triplicates) were labeled using TMT 6-plex reagents, with control samples assigned to reporter ions 126, 127, 128 and treated samples to 129, 130, 131. Labeling efficiency was verified before pooling equal amounts of peptides across channels. The combined peptide mixture was vacuum-dried and subjected to phosphopeptide enrichment using Fe-NTA affinity chromatography according to the manufacturer’s instructions. Enriched phosphopeptides were analyzed on an Orbitrap Fusion Tribrid mass spectrometer coupled to an EASY-nLC 1200 system. Peptides were separated on a C18 nano-column using a linear gradient of 5–35% acetonitrile (0.1% formic acid) over 90 min at a flow rate of 300 nL/min. MS data were acquired in data-dependent acquisition mode with full MS scans collected at 120,000 resolution (m/z 400–1600), followed by HCD fragmentation of precursor ions (charge states 2–6) using a top-speed method (3 s cycle time). MS/MS spectra were acquired at 30,000 resolution with dynamic exclusion set to 30 s.

Raw files were processed using Proteome Discoverer (v2.2) and searched against the human RefSeq database using SEQUEST and Mascot algorithms. Static modifications included carbamidomethylation of cysteine and TMT labeling at peptide N-termini and lysine residues, while phosphorylation (Ser/Thr/Tyr) and methionine oxidation were set as variable modifications. Precursor and fragment mass tolerances were set to 10 ppm and 0.05 Da, respectively, with a 1% false discovery rate at the peptide-spectrum match level. Differential phosphorylation analysis was performed using Perseus, and functional enrichment analyses were conducted using DAVID and Reactome pathway databases.

### Molecular docking of metabolites with target proteins

The crystallographic 3D structures of proteins dUTP, GMPR, POLA1, and CPS1 were obtained from RCSB-PDB and the 3D crystallographic structure of protein UCKL1 was retrieved from AlphaFold Protein Structure Database. Protein structures were prepared using the Protein Preparation Wizard in Schrödinger-2025-4, which involved the addition of missing hydrogen atoms, assignment of bond orders, optimization of hydrogen bonding networks, and removal of water molecules beyond 8 Å from hetero groups using PROPKA and Epik 2.0. Protonation states of residues were assigned at physiological pH (7.4), and energy minimization was performed using the OPLS4 force field. Metabolite structures, including deoxyuridine triphosphate, guanosine monophosphate, and uridine monophosphate, were retrieved from the PubChem. Ligands were prepared using the LigPrep module, where ionization states were generated at pH 7.0 ± 2.0, stereochemistry was retained, and energy minimization was performed using the OPLS4 force field. The structures were desalted and tautomers were generated. Top ranked potentioal active binding sites were generated. Followed by receptor grid generation by defining the active site based on co-crystallized ligands or predicted binding pockets.

Molecular docking analysis was performed using the Schrödinger Suite (Schrödinger-2025-4, LLC, New York, USA) to evaluate potential interactions between significantly altered metabolites and target proteins identified from proteomic analysis.Three-dimensional structures of target proteins, including DUT, GMPR, and CPS1, were obtained from the Protein Data Bank. Protein structures were prepared using the Protein Preparation Wizard in Schrödinger, which involved the addition of missing hydrogen atoms, assignment of bond orders, optimization of hydrogen bonding networks, and removal of water molecules beyond 5 Å from hetero groups. Protonation states of residues were assigned at physiological pH (7.0), and energy minimization was performed using the OPLS4 force field.

Total of 13 dysregulated metabolite structures were retrieved from the PubChem in .sdf format. Ligands were prepared using the LigPrep module, where ionization states were generated at pH 7.0 ± 0.5, stereochemistry was retained, and energy minimization was performed using the OPLS4 force field.Receptor grids were generated by defining the active site based on co-crystallized ligands or predicted binding pockets. Molecular docking was performed using the Glide module in Schrödinger-2025-4 in standard precision (SP) mode. Default parameters were used, and flexible ligand docking was enabled to allow conformational sampling of ligands within the binding site.

Docking poses were ranked based on Glide docking scores, and the best-scoring pose for each ligand–protein pair was selected for further analysis.Protein–ligand interactions were analyzed using Maestro visualization tools to identify hydrogen bonds, hydrophobic interactions, salt bridges, and π–π stacking interactions. Two-dimensional interaction diagrams were generated to visualize binding modes and key interacting residues.

## RESULTS

Cancer cells undergo profound metabolic reprogramming to sustain rapid proliferation, biomass accumulation, and survival [31]. Although CAMKK2 is overexpressed in multiple malignancies, including gastric cancer, its role in regulating cancer-associated metabolic pathways remains incompletely understood. Given the established involvement of CAMKK2 in cellular energy sensing and kinase signaling networks, we hypothesized that CAMKK2 may function as a key metabolic regulator in gastric cancer.

To investigate this, gastric cancer cell lines (AGS) were treated with the selective CAMKK2 inhibitor STO-609 and subjected to comprehensive untargeted metabolomic profiling. To mechanistically link metabolic alterations with upstream signaling events, we further performed integrated phosphoproteomic and quantitative proteomic analyses. This multi-omics approach enabled systematic evaluation of how CAMKK2 inhibition reshapes cellular metabolism and how these metabolic changes are coordinated with alterations in kinase signaling and protein expression programs that govern cell cycle progression and DNA replication.

### CAMKK2 inhibition induces global metabolic reprogramming in gastric cancer cells

To investigate the metabolic consequences of CAMKK2 inhibition in gastric cancer, AGS cells were treated with the selective CAMKK2 inhibitor STO-609 and subjected to LC–MS/MS–based untargeted metabolomic profiling in both positive and negative ionization modes. The overall experimental workflow, including metabolite extraction, chromatographic separation, and high-resolution mass spectrometry–based detection, is illustrated in Figure 1A.

**Figure 1.**
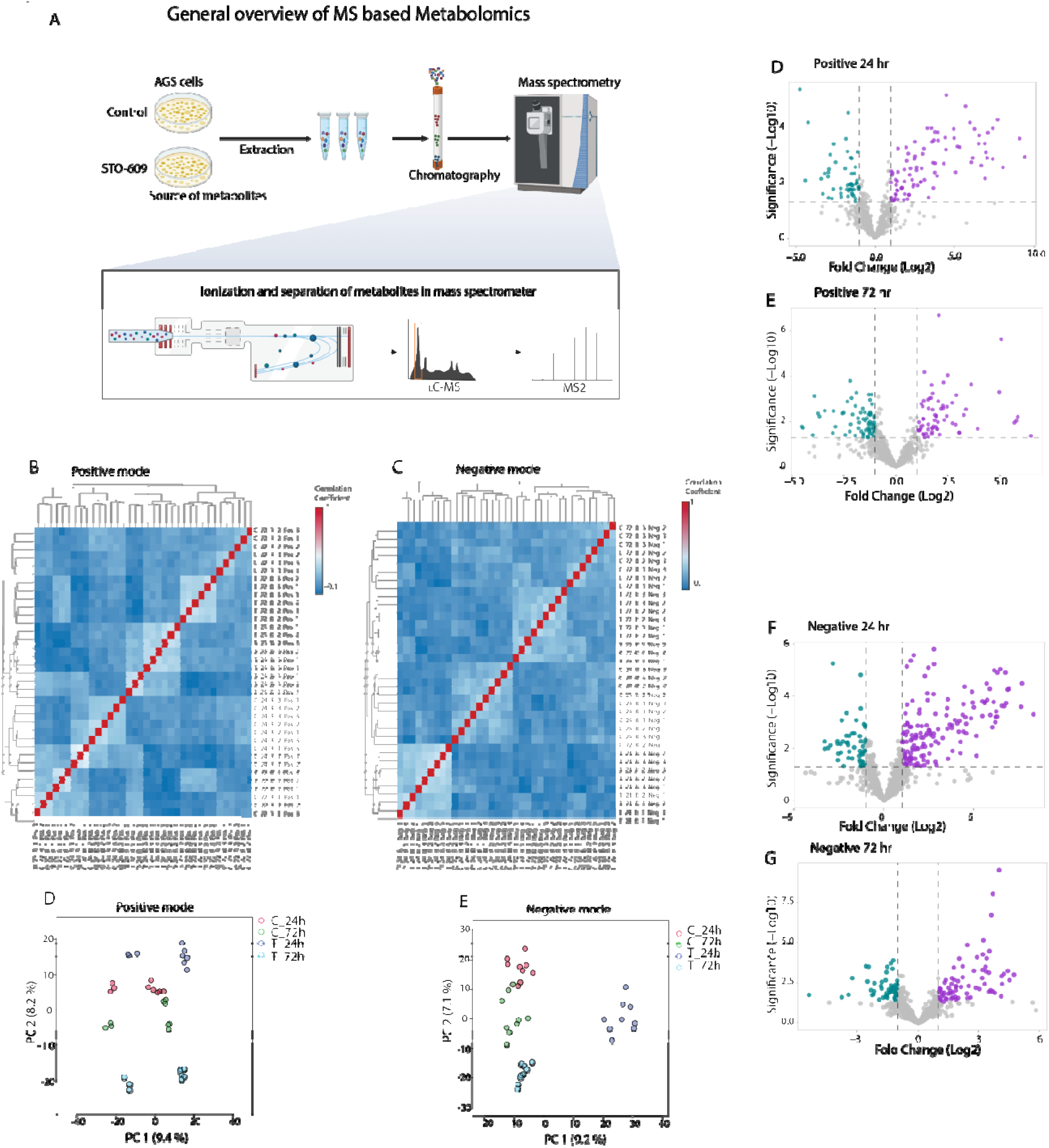
CAMKK2 inhibition induces global metabolic reprogramming in gastric cancer cells: (A) Schematic overview of the untargeted LC–MS/MS-based metabolomics workflow, including metabolite extraction, chromatographic separation, and mass spectrometry analysis. (B–C) Pearson correlation heatmaps showing reproducibility and clustering of biological replicates in positive (B) and negative (C) ionization modes. (D–E) Principal component analysis (PCA) plots demonstrating separation between control and STO-609-treated samples at 24 h and 72 h in positive (D) and negative (E) modes. (F–G) Volcano plots depicting significantly altere metabolic features at 24 h and 72 h following CAMKK2 inhibition in positive (F) and negative (G) ionization modes. Significance thresholds were defined based on fold change and statistical criteria.

Following data preprocessing, including peak detection, alignment, normalization, and filtering, a total of positive 1,5799 and negative 1,8605 reproducible metabolic features were identified across all biological replicates detailed information listed in Table 1. Quality assessment demonstrated high analytical reproducibility, with consistent signal intensity distributions and retention time alignment across samples. Pearson correlation analysis revealed moderate to strong clustering of biological replicates within groups in both positive and negative ionization modes (Figure 1B-C), indicating reliable data acquisition while also capturing biologically relevant variability induced by CAMKK2 inhibition. Importantly, no major outliers were observed, supporting the robustness and integrity of the dataset.

**Table 1:** Distribution of metabolites across positive and negative ionization modes, including features, MS1 detection, MS2 annotation, and human metabolite mapping.

| Mode | Features | MS1 | MS2 | Human mapped |
| --- | --- | --- | --- | --- |
| Positive | 1,5799 | 879 | 365 | 45 |
| Negative | 1,8605 | 988 | 344 | 61 |

To further evaluate global metabolic variation, principal component analysis (PCA) was performed separately for positive and negative ionization datasets. PCA revealed clear and distinct separation between control and CAMKK2-inhibited samples across both ionization modes (Figure 1D-E), demonstrating a pronounced metabolic shift upon inhibition of CAMKK2. Notably, samples also exhibited time-dependent clustering, with distinct grouping of 24 h and 72 h samples, suggesting progressive and dynamic metabolic remodeling over time.

To identify significantly altered metabolites, differential analysis was performed using fold-change and statistical significance thresholds. Volcano plot analysis revealed widespread metabolic perturbations at both 24 h and 72 h time points (Figure 1F–G), with a substantial number of features meeting significance criteria. A large proportion of significantly altered metabolites were downregulated, indicating suppression of key metabolic pathways following CAMKK2 inhibition. In contrast, a subset of metabolites exhibited upregulation, suggesting potential compensatory metabolic responses during early stages of inhibition.

To further enhance confidence in metabolite identification, MS/MS (MS2)-based annotation was performed, and confidently matched metabolites were subjected to hierarchical clustering analysis. Heatmaps of MS2-validated metabolites demonstrated clear segregation between control and treated samples in both positive and negative ionization modes (Supplementary Figure 1A-B). These clustering patterns further corroborate the reproducibility and biological relevance of the observed metabolic alterations, while also confirming the robustness of metabolite annotation.

Collectively, these results demonstrate that pharmacological inhibition of CAMKK2 induces broad, reproducible, and time-dependent metabolic reprogramming in gastric cancer cells. The observed global suppression of metabolite abundance, combined with validated metabolite identification, supports a central role for CAMKK2 in maintaining metabolic homeostasis and highlights its importance in regulating cancer cell metabolism.

### CAMKK2 inhibition induces time-dependent metabolic rewiring with suppression of nucleotide and lipid metabolic pathways

To further characterize the metabolic alterations induced by CAMKK2 inhibition, significantly altered metabolites were subjected to hierarchical clustering and pathway enrichment analyses. Heatmaps of differentially abundant metabolites revealed clear and consistent separation between control and STO-609-treated samples across both time points and ionization modes.

At 24 hours, hierarchical clustering demonstrated distinct metabolic signatures between control and treated groups in both positive and negative ionization modes (Figure 2A-B). A subset of metabolites exhibited increased abundance in treated samples, whereas a larger proportion showed reduced levels, suggesting early metabolic perturbations following CAMKK2 inhibition. At 72 hours, these differences became more pronounced, with stronger segregation of control and treated samples observed in both positive and negative modes (Figure 2C-D). Notably, a widespread reduction in metabolite abundance was evident at this later time point, indicating progressive metabolic suppression upon sustained CAMKK2 inhibition.

**Figure 2.**
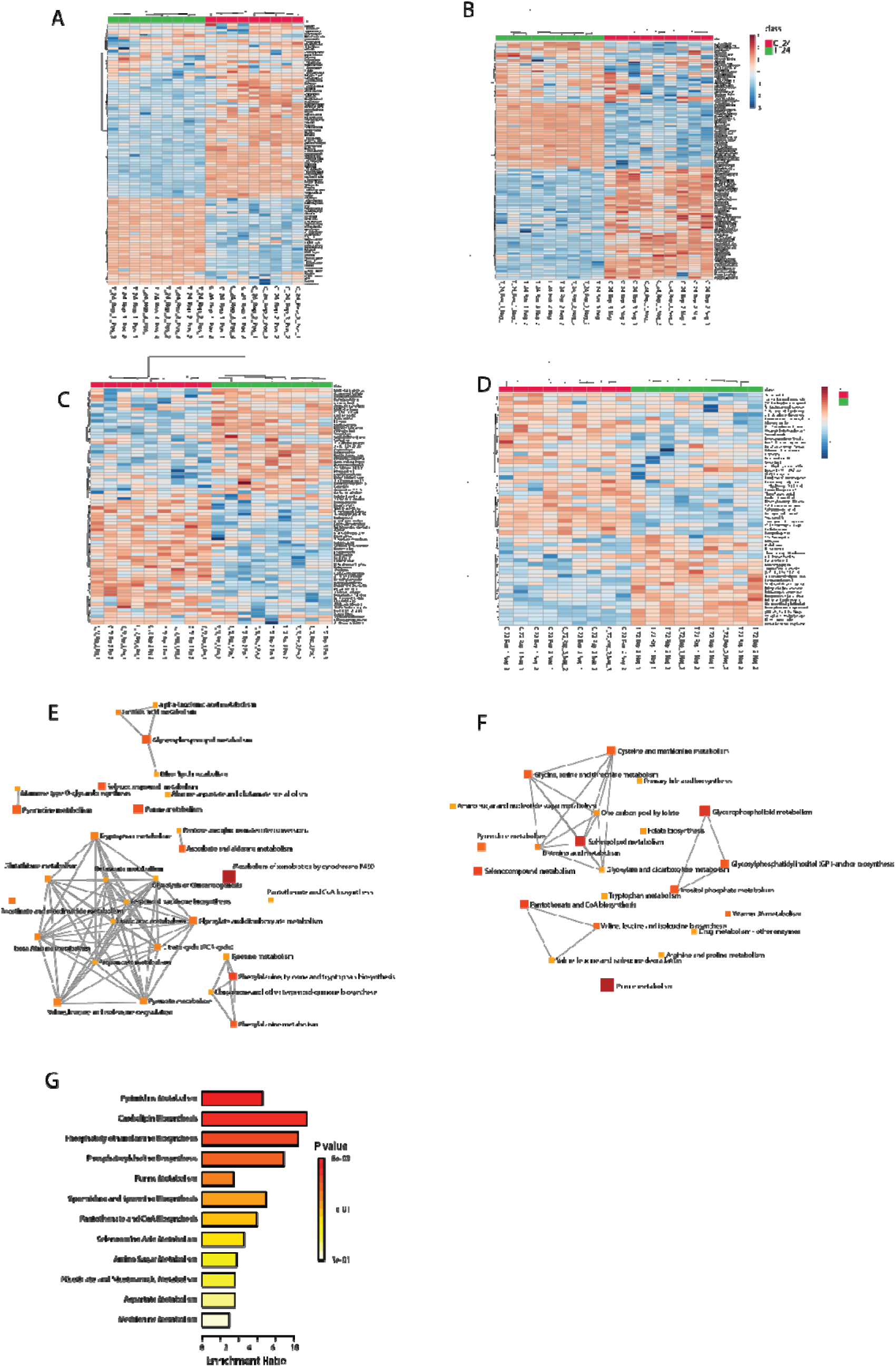
CAMKK2 inhibition induces time-dependent metabolic rewiring in gastric cancer cells: (A–B) Heatmaps of significantly altered metabolites at 24 hours post-treatment in positive (A) and negative (B) ionization modes, showing distinct clustering between control and STO-609-treated samples. (C–D) Heatmaps of significantly altered metabolites at 72 hours post-treatment in positive (C) and negative (D) ionization modes, demonstrating enhanced separation and progressive metabolic changes over time. (E) Pathway enrichment analysis of all identified metabolites in positive ionization mode, highlighting key metabolic pathways altered upon CAMKK2 inhibition. (G) Pathway enrichment analysis of downregulated metabolites following CAMKK2 inhibition, showing significant enrichment of nucleotide metabolism, phospholipid metabolism, and central metabolic pathways. Enrichment significance is represented by color intensity and enrichment ratio.

To gain functional insights into these metabolic changes, pathway enrichment analysis was performed using significantly altered metabolites. In the positive ionization mode, enriched pathways included amino acid metabolism, purine metabolism, pyrimidine metabolism, and lipid metabolic pathways, highlighting broad metabolic remodeling (Figure 2E). These pathways are critically associated with cellular proliferation and biosynthetic processes. Further focused analysis of downregulated metabolites revealed significant enrichment of pathways related to nucleotide metabolism, phospholipid metabolism, and central carbon metabolism, including purine metabolism, phosphatidylcholine biosynthesis, and glycolysis-related pathways (Figure 2G). The consistent downregulation of these pathways suggests impaired nucleotide biosynthesis and membrane lipid production, which are essential for DNA replication and cell proliferation.

To validate pathway-level interpretations, enrichment analysis was also performed using MS/MS (MS2)-confirmed metabolites. Enrichment networks generated from MS2-annotated metabolites in both positive and negative ionization modes demonstrated similar pathway distributions, further supporting the robustness of metabolite identification (Supplementary Figure 2A–B).

In addition, pathway analysis restricted to metabolites mapped to the human metabolic database revealed consistent enrichment patterns, particularly in pathways related to amino acid metabolism, nucleotide metabolism, and mitochondrial energy metabolism in both ionization modes (Supplementary Figure 2C-D). These findings reinforce the biological relevance of the observed metabolic alterations.

Collectively, these results demonstrate that CAMKK2 inhibition leads to time-dependent metabolic rewiring, characterized by progressive suppression of nucleotide, lipid, and central metabolic pathways. This metabolic shift is consistent with reduced biosynthetic capacity and aligns with the observed inhibition of DNA replication and cell cycle progression identified in proteomic analyses.

### CAMKK2 inhibition suppresses cell cycle progression and induces replication stress through coordinated proteomic and metabolic remodeling

To investigate the molecular consequences of CAMKK2 inhibition, we integrated quantitative proteomics with metabolomic datasets to identify key pathways and regulatory networks affected by CAMKK2 signaling. Unsupervised clustering of differentially expressed proteins revealed a distinct subset of proteins that were consistently downregulated upon CAMKK2 inhibition (Figure 3A). These proteins were predominantly associated with DNA replication, cell cycle progression, chromatin organization, and DNA repair, indicating a coordinated suppression of proliferative machinery. A comprehensive heatmap of downregulated proteins further confirmed this global trend (Supplementary Figure 3A). These findings are consistent with previous studies demonstrating that disruption of replication-associated proteins and chromatin regulators leads to impaired S-phase progression and proliferative arrest [40, 48].

**Figure 3.**
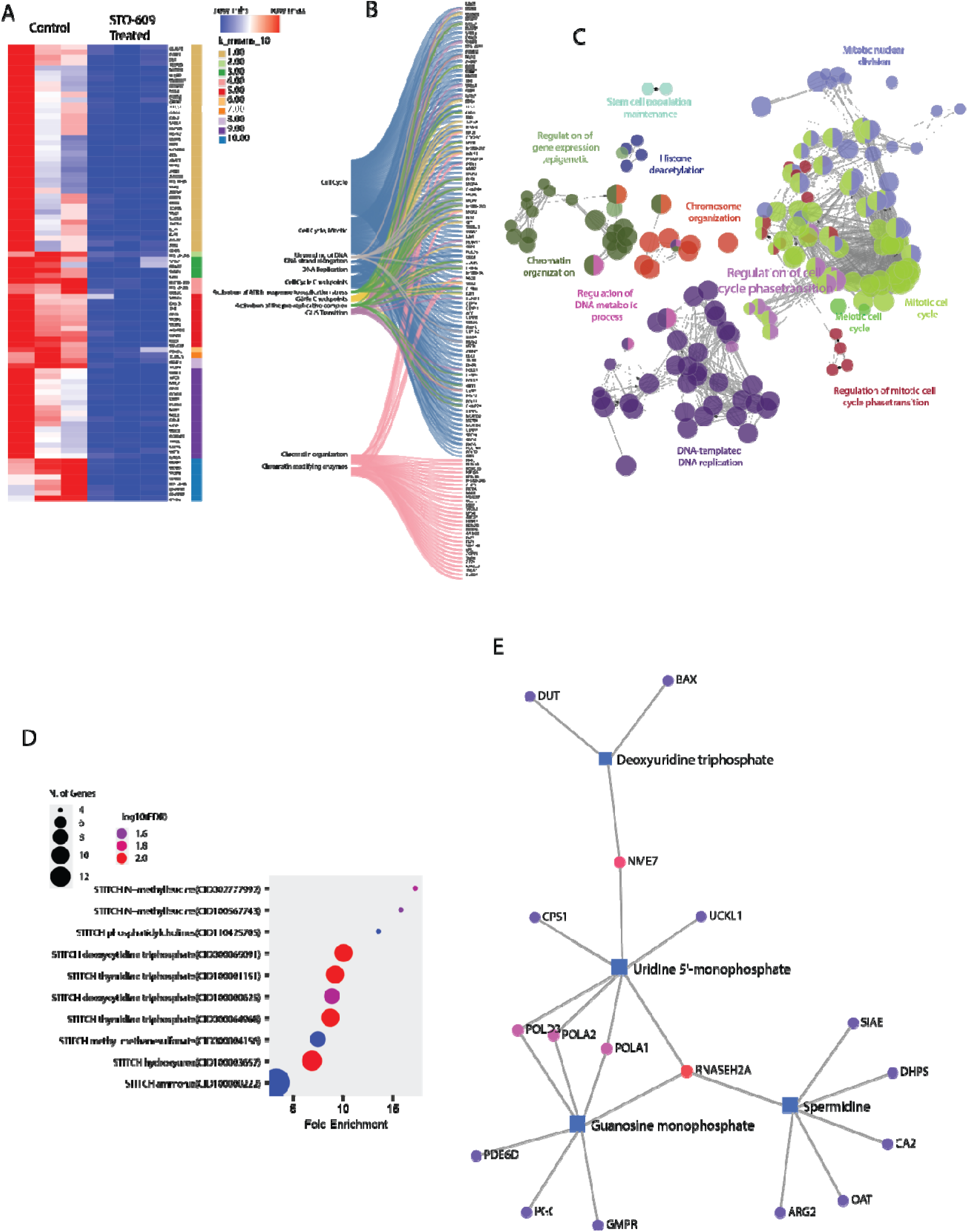
Integrated proteomic and metabolomic analysis reveals suppression of cell cycle progression and nucleotide metabolism upon CAMKK2 inhibition: (A) Heatmap of differentially expressed proteins showing a distinct cluster of downregulated proteins associated with DNA replication, cell cycle progression, chromatin organization, and DNA repair in STO-609-treated samples compared to controls. (B) Reactome pathway enrichment analysis of downregulated proteins, highlighting significant enrichment of pathways related to DNA replication, cell cycle regulation, DNA damage response, and chromatin remodeling. (C) Protein–protein interaction network (BioGRID) illustrating highly interconnected clusters of proteins involved in mitotic cell cycle regulation, DNA-templated replication, and chromosomal organization. (D) Functional enrichment analysis (ShinyGO v0.85.1 and Drug GeneSetDB) showing overrepresentation of pathways associated with DNA replication, chromosome segregation, and replication stress–related perturbations, including signatures linked to nucleotide metabolism inhibitors. (E) Integrated protein–metabolite interaction network linking downregulated metabolites (including deoxyuridine triphosphate, uridine monophosphate, and guanosine monophosphate) with key enzymes involved in nucleotide metabolism and DNA replication (e.g., DUT, GMPR, POLA1, POLA2), highlighting coordinated disruption of metabolic and replication machinery.

To further characterize the functional implications of these changes, pathway enrichment analysis was performed. Reactome-based analysis demonstrated that downregulated proteins were enriched in pathways related to cell cycle regulation, DNA synthesis, DNA damage response, and chromatin remodeling (Figure 3B). These pathways converge on key biological processes governing DNA replication, chromosome segregation, and mitotic progression, which are tightly regulated by kinase-driven signaling networks [49, 50]. The coordinated suppression of these pathways suggests that CAMKK2 plays a critical role in sustaining proliferative signaling required for DNA replication and mitosis. Protein–protein interaction analysis using BioGRID revealed highly interconnected clusters of proteins involved in mitotic cell cycle regulation, DNA-templated replication, and chromosomal organization (Figure 3C). These tightly connected modules are characteristic of replication-associated protein networks. Disruption of such networks has been shown to impair replication fork progression and increase genomic instability.

Gene ontology enrichment analysis performed using ShinyGO (v0.85.1) further supported these findings. Biological process enrichment analysis demonstrated significant overrepresentation of pathways related to DNA replication, mitotic nuclear division, chromosome segregation, and cell cycle regulation (Supplementary Figure 3B). Consistently, molecular function and cellular component analyses revealed enrichment of DNA binding, helicase activity, replication fork components, and chromosomal complexes (Figure 3D and Supplementary data). These functional categories are essential for replication origin firing and fork stability, processes that are highly sensitive to disruptions in signaling and metabolic inputs [51, 52].

To explore functional associations with pharmacological perturbations, enrichment analysis using Drug GeneSetDB was performed. This analysis revealed significant enrichment of gene sets associated with nucleotide metabolism inhibitors and replication stress–inducing agents, including signatures related to hydroxyurea and nucleotide analog mediated perturbations (Figure 3D). Hydroxyurea, a well-established inhibitor of ribonucleotide reductase, induces replication stress by depleting intracellular deoxyribonucleotide pools required for DNA synthesis [51, 52]. The similarity between these signatures and CAMKK2 inhibition suggests that loss of CAMKK2 activity induces a cellular state resembling nucleotide insufficiency–driven replication stress.

To establish a direct link between proteomic alterations and metabolic rewiring, integrated protein–metabolite interaction analysis was performed. Network analysis demonstrated that several downregulated metabolites, including deoxyuridine triphosphate, uridine monophosphate, and guanosine monophosphate, were functionally connected to key enzymes involved in nucleotide metabolism and DNA replication (Figure 3E). Notably, proteins such as DUT, GMPR, POLA1, and POLA2, which play essential roles in nucleotide biosynthesis and DNA replication, were linked to altered metabolite levels. This observation is consistent with previous reports demonstrating that depletion of nucleotide pools directly impairs DNA polymerase activity and replication fork progression, thereby inducing replication stress [40, 53]. Collectively, these findings demonstrate that CAMKK2 inhibition induces a coordinated suppression of cell cycle progression, DNA replication machinery, and nucleotide metabolism, resulting in a cellular phenotype consistent with replication stress. This multi-layered disruption highlights a critical role for CAMKK2 in integrating signaling and metabolic pathways to maintain proliferative capacity and genomic stability in gastric cancer cells.

### CAMKK2 inhibition remodels phosphorylation-dependent signaling networks linking cell cycle regulation to metabolic reprogramming

To further delineate the signaling mechanisms underlying CAMKK2-mediated metabolic regulation, we integrated phosphoproteomics with metabolomics datasets to identify phosphorylation-dependent pathways affected by CAMKK2 inhibition. Global phosphoproteomic analysis revealed widespread alterations in phosphorylation upon CAMKK2 inhibition. Unsupervised clustering identified a distinct subset of hypophosphorylated peptides corresponding to proteins involved in DNA replication, cell cycle progression, and chromatin organization (Figure 4A). A comprehensive heatmap of hypophosphorylated peptides further confirmed this global trend (Supplementary Figure 4A), indicating that CAMKK2 activity is critical for maintaining phosphorylation-dependent regulation of proliferative pathways. These observations are consistent with previous reports demonstrating that phosphorylation events tightly regulate replication origin firing, chromatin accessibility, and mitotic progression [54–56]. To characterize the functional implications of these phosphorylation changes, pathway enrichment analysis was performed. Sankey plot–based visualization demonstrated that hypophosphorylated proteins were enriched in pathways associated with DNA damage response, DNA repair, chromatin remodeling, DNA replication, and cell cycle regulation (Figure 4B). These pathways converge on essential biological processes governing DNA synthesis and mitotic progression, which are known to be highly dependent on kinase-mediated phosphorylation cascades [57, 58]. The observed suppression of these pathways suggests that CAMKK2 plays a key role in sustaining phosphorylation networks required for cell cycle progression.

**Figure 4.**
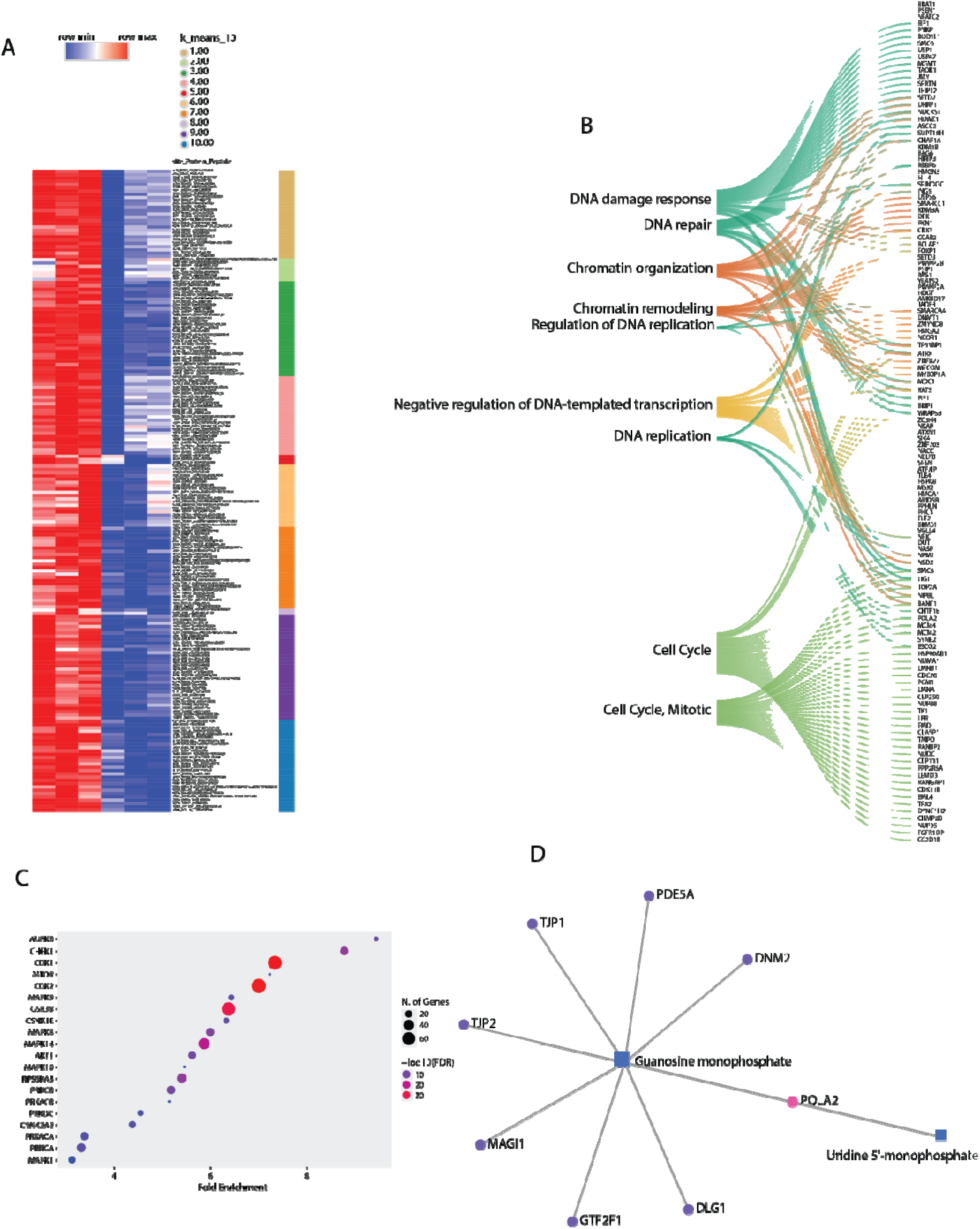
Phosphoproteomic analysis reveals disruption of kinase signaling and coupling to nucleotide metabolism upon CAMKK2 inhibition: (A) Heatmap of hypophosphorylated peptides in STO-609-treate samples compared to controls, highlighting proteins involved in DNA replication, cell cycle progression, an chromatin organization. (B) Sankey plot–based pathway enrichment analysis of hypophosphorylated proteins, showing enrichment of pathways associated with DNA damage response, DNA repair, chromatin remodeling, DNA replication, and cell cycle regulation. (C) Kinase enrichment analysis (KEA 2015 via ShinyGO v0.85.1) identifying upstream kinases associated with the observed phosphorylation changes, including CDK family kinases, MAPK signaling components, and checkpoint regulators. (D) Integrated protein–metabolite interaction network linking hypophosphorylated proteins with downregulated metabolites involved in nucleotide metabolism, including guanosine monophosphate and uridine monophosphate, highlighting functional coupling between signaling pathways and metabolic regulation.

To identify upstream regulatory kinases associated with these phosphorylation changes, kinase enrichment analysis was performed using KEA (KEA 2015) via ShinyGO (v0.85.1). This analysis revealed significant enrichment of kinases involved in cell cycle control and stress signaling, including CDK family kinases, MAPK signaling components, and checkpoint regulators (Figure 4C). CDKs are essential drivers of cell cycle transitions and replication initiation, while MAPK pathways regulate proliferation and stress responses [58, 59]. The coordinated attenuation of these kinase networks upon CAMKK2 inhibition indicates disruption of signaling cascades that orchestrate DNA replication and mitotic progression.

To establish a functional connection between phosphorylation-dependent signaling and metabolic alterations, integrated protein-metabolite interaction analysis was performed. Network analysis revealed that hypophosphorylated proteins were functionally linked to downregulated metabolites involved in nucleotide metabolism, including guanosine monophosphate and uridine monophosphate (Figure 4D). Notably, proteins such as POLA2, which play essential roles in DNA replication, were directly associated with altered metabolite levels. This observation is consistent with emerging evidence that kinase signaling pathways are tightly coupled to metabolic processes to ensure adequate nucleotide supply for DNA synthesis [60]Gene ontology enrichment analysis of hypophosphorylated proteins further supported these findings. Biological process enrichment analysis demonstrated significant overrepresentation of pathways related to RNA processing, chromatin organization, DNA metabolic processes, and regulation of biosynthetic pathways (Supplementary Figure 4B). Cellular component analysis revealed enrichment in nuclear compartments, chromatin-associated complexes, and replication-associated structures, while molecular function analysis highlighted DNA binding, chromatin binding, and protein interaction activities (Supplementary Figure 4C–D).

Collectively, these results demonstrate that CAMKK2 inhibition leads to widespread remodeling of phosphorylation-dependent signaling networks, resulting in suppression of cell cycle progression, DNA replication, and chromatin regulation. Integration with metabolomic data further reveals a direct link between disrupted signaling pathways and nucleotide metabolic insufficiency, reinforcing a model in which CAMKK2 coordinates signaling and metabolic programs to sustain proliferative capacity in gastric cancer cells.

### Integrated docking and multi-omics analysis reveals disruption of metabolite–enzyme coupling following CAMKK2 inhibition

To further investigate the mechanistic relationship between altered metabolite levels and enzyme function, molecular docking analysis was performed between significantly dysregulated nucleotide metabolites and key proteins identified from proteomic analysis. Docking analysis revealed favorable binding interactions between nucleotide metabolites and enzymes involved in nucleotide metabolism and DNA replication. Specifically, deoxyuridine triphosphate exhibited strong binding affinity toward DUT (dUTPase) with a docking score of −10.7, indicating a stable interaction within the catalytic pocket (Figure 5A). Similarly, guanosine monophosphate showed robust binding to GMPR with a docking score of −11.5 (Figure 5B), while uridine monophosphate demonstrated favorable interaction with CPS1 (score −9) (Figure 5C). These interactions were supported by multiple hydrogen bonds, polar contacts, and hydrophobic interactions within the binding sites, suggesting structurally plausible metabolite enzyme associations.

**Figure 5.**
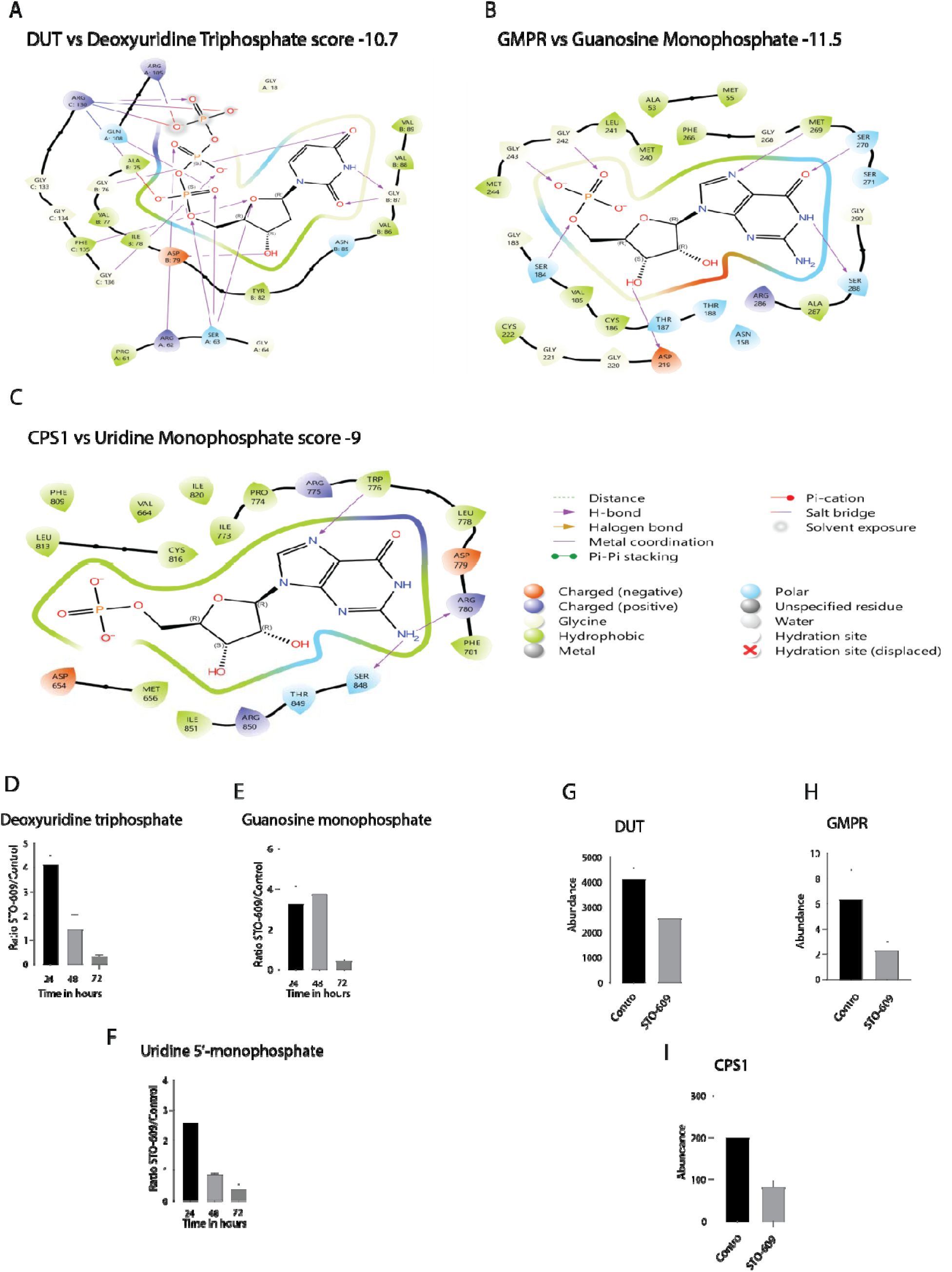
CAMKK2 inhibition disrupts nucleotide metabolism through altered metabolite–enzyme interactions and temporal metabolic depletion: (A-C) Molecular docking analysis showing binding interactions between nucleotide metabolites and key enzymes: deoxyuridine triphosphate with DUT (A), guanosine monophosphate with GMPR (B), and uridine monophosphate with CPS1 (C). Docking scores indicate favorable binding affinities, with interactions stabilized by hydrogen bonds, polar contacts, and hydrophobic interactions within the catalytic pockets. (D–F) Temporal analysis of metabolite abundance at 24 h, 48 h, and 72 h following CAMKK2 inhibition, showing an early increase followed by progressive depletion of nucleotide metabolites. (G–I) Protein abundance of corresponding enzymes (DUT, GMPR, CPS1) in control and STO-609-treated samples at 72 h, demonstrating reduced expression upon CAMKK2 inhibition.

Detailed characterization of the active site residues responsible for these interactions is presented in Table 2. At the atomic level, the binding of deoxyuridine triphosphate within the DUT catalytic pocket was stabilized by hydrogen bonds formed with residues SER A:63, ARG A:62, GLY C: 136, PHE C:135, GLY B:76, GLN A:108, ARG C:130, GLY B: 87, and ASP B:79, Salt bridges such as ARG C:130, and ARG A:105, spanning all three protein chains (A,B, and C) consistent with the structure of dUTPase in which the active site is assembled at the subunit interference. For GMPR, the binding of guanosine monophosphate was mediated by hydrogen bonds with residues GLY: 242, GLY:243, SER: 184, ASP:219, SER: 288, SER: 270, and MET: 269, with no salt bridge or aromatic contacts detected, indicating that hydrogen bonding is the primary driver of selectivity within a hydrophilic active site pocket. In case of CPS1, uridine monophosphate engaged hydrogen bonds with TRP:776, and LEU:778, and further stabilized by pi-cation interaction with ARG:775 and pi-pi stacking with PHE:781, suggesting that an aromatic hydrophobic microenvironment within the CPS1 binding pocket. Lastly, UCKL1 bound uridine monophosphate through hydrogen bonds with ASN:201, and PRO:86 residues.

**Table 2:** Molecular docking analysis of the four selected target proteins against their respective ligand revealed favorable binding interactions.

| Protein | Ligand | Docking score | Active site residues and types of bond formation |
| --- | --- | --- | --- |
| DUT<br>(dUTPase) | deoxyuridine triphosphate | -10.7 | HBonds: SER A:63, ARG A:62, GLY C: 136, PHE C:135, GLY B:76, GLN A:108, ARG C:130, GLY B: 87, and ASP B:79 |
| GMPR | guanosine monophosphate | -11.5 | HBonds: GLY: 242, GLY:243, SER: 184, ASP:219, SER: 288, SER: 270, and MET: 269 |
| CPS1 | Uridine monophosphate | -9.0 | HBonds: TRP:776, and LEU:778<br>Pi-cation: ARG:775<br>Pi-Pi- stacking: PHE:781 |
| UCKL1 | Uridine Monophosphate | -7.1 | HBonds: ASN:201, and PRO:86 |

To assess the temporal dynamics of these metabolites following CAMKK2 inhibition, their relative abundance was examined at 24, 48, and 72 hours. Notably, deoxyuridine triphosphate levels showed an initial increase at 24 hours, followed by a marked decline at later time points (Figure 5D). A similar temporal pattern was observed for guanosine monophosphate and uridine monophosphate, with early elevation or maintenance followed by significant reduction at 48 and 72 hours (Figure 5E–F). This trend suggests an early compensatory response in nucleotide pools, followed by progressive depletion consistent with impaired nucleotide biosynthesis.

Consistent with these metabolite changes, proteomic analysis revealed significant downregulation of the corresponding enzymes at 72 hours following CAMKK2 inhibition. Specifically, DUT, GMPR, and CPS1 protein levels were reduced compared to control conditions (Figure 5G-I), further supporting disruption of nucleotide metabolic pathways at the protein level. Importantly, the integration of docking, metabolite abundance, and proteomic data highlights a coordinated disruption of metabolite–enzyme coupling. While docking analysis supports the structural feasibility of interactions between nucleotide metabolites and their respective enzymes, the observed reduction in both metabolite levels and enzyme abundance at later time points suggests impaired catalytic flux through nucleotide biosynthesis pathways.

Collectively, these findings provide multi-layered evidence that CAMKK2 inhibition disrupts nucleotide metabolism through both structural and functional mechanisms, ultimately leading to depletion of nucleotide pools and suppression of DNA replication. This integrative analysis reinforces a model in which CAMKK2 regulates metabolic and enzymatic networks required for maintaining proliferative capacity in gastric cancer cells.

### CAMKK2 inhibition functionally validates multi-omics findings by inducing mitotic defects and multinucleation

To functionally validate the multi-omics observations indicating suppression of DNA replication, cell cycle progression, and nucleotide metabolism, we examined the impact of CAMKK2 inhibition on nuclear morphology and cell division using DAPI staining. AGS cells treated with STO-609 exhibited progressive and pronounced alterations in nuclear architecture over time (24-96 h) (Figure 6A). In control cells, nuclei were uniformly shaped and predominantly mononucleated, consistent with normal proliferative capacity. In contrast, CAMKK2-inhibited cells displayed a time-dependent reduction in cell number, indicative of impaired proliferation (Figure 6A-B).

**Figure 6.**
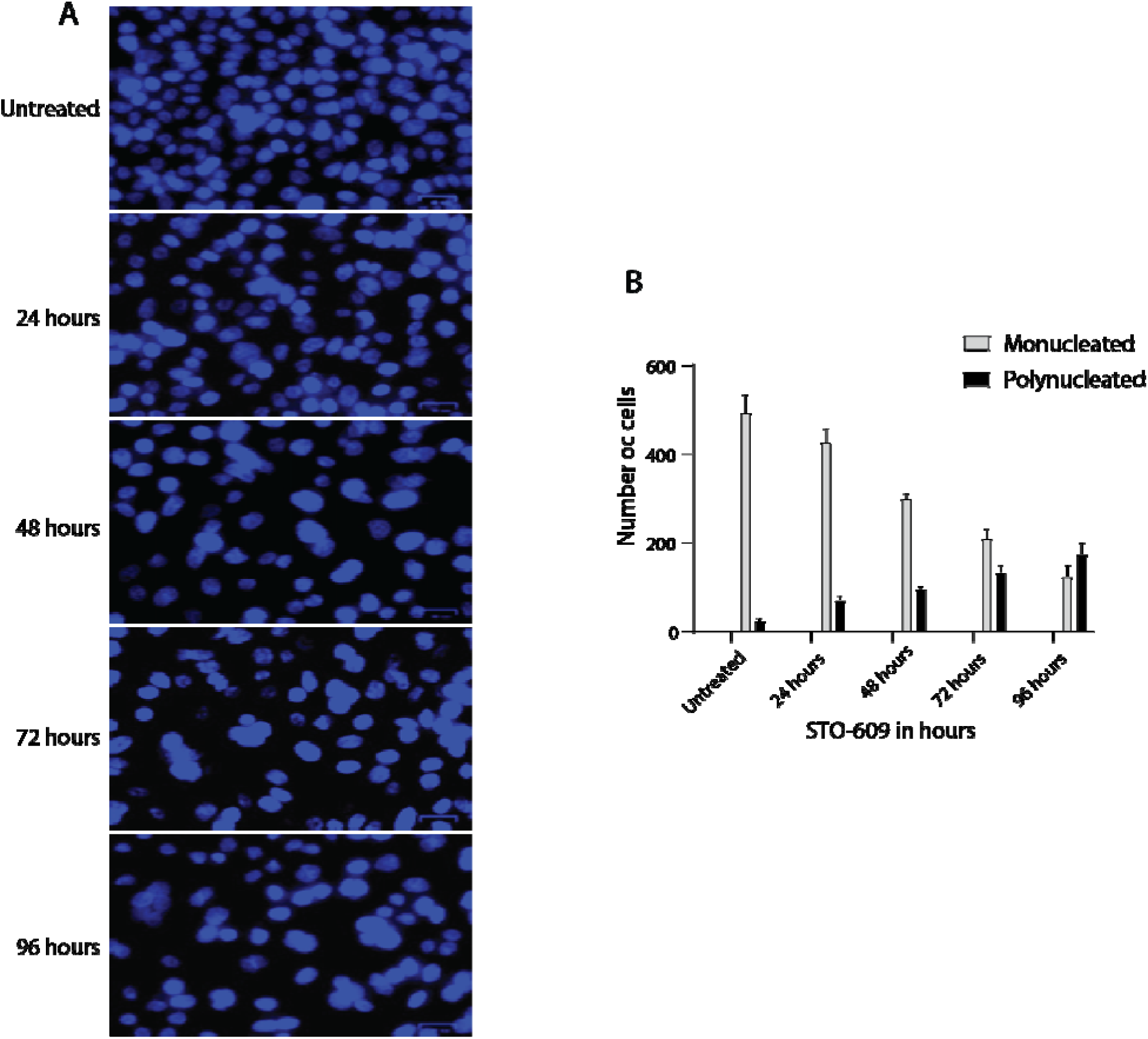
CAMKK2 inhibition induces multinucleation and reduced cell proliferation: (A) Representative DAPI-stained images of AGS cells treated with STO-609 at indicated time points show decreased cell density and increased multinucleation over time. (B) Quantification of mononucleated and multinucleated cells demonstrates a time-dependent decrease in mononucleated cells and increase in multinucleated cells following CAMKK2 inhibition.

Strikingly, CAMKK2 inhibition led to a progressive accumulation of multinucleated cells, accompanied by enlarged and irregular nuclear morphology (Figure 6A). These features are hallmarks of mitotic defects and failed cytokinesis, suggesting that cells are unable to successfully complete cell division. Quantitative analysis confirmed a significant shift in nuclear states (Figure 6B), with a marked decrease in mononucleated cells and a corresponding increase in multinucleated cells at later time points. This phenotype is consistent with replication stress and defective mitotic progression, as cells attempt to divide in the context of insufficient nucleotide pools and impaired replication machinery. Importantly, these phenotypic changes directly align with our multi-omics findings. The observed multinucleation and proliferation defects correlate with (i) downregulation of DNA replication and cell cycle proteins (Figure 3), (ii) suppression of phosphorylation-dependent signaling pathways governing mitotic progression (Figure 4), and (iii) depletion of key nucleotide metabolites (Figures 1–2). Together, these data support a model in which CAMKK2 coordinates metabolic and signaling networks required for proper DNA synthesis and cell division.

Collectively, these results provide functional validation that CAMKK2 inhibition disrupts proliferative capacity by inducing replication stress, mitotic failure, and genomic instability, thereby reinforcing the central role of CAMKK2 in maintaining cell cycle fidelity in gastric cancer cells.

### Integrated multi-omics analysis defines a CAMKK2-dependent regulatory axis linking kinase signaling, nucleotide metabolism, and cell cycle progression

To synthesize the multi-layered effects of CAMKK2 inhibition, we developed an integrative model summarizing the coordinated molecular and phenotypic alterations observed across metabolomic, proteomic, phosphoproteomic, and functional analyses (**Figure 7**). CAMKK2 inhibition resulted in a marked suppression of kinase-driven signaling networks, as evidenced by reduced phosphorylation of CDK, MAPK, and mitotic kinase targets. This global hypophosphorylation was associated with disruption of key regulatory pathways governing DNA replication and cell cycle progression. Consistently, proteomic analysis revealed downregulation of critical replication-associated proteins, including components of the MCM complex, DNA polymerases, and cell cycle regulators.

**Figure 7.**
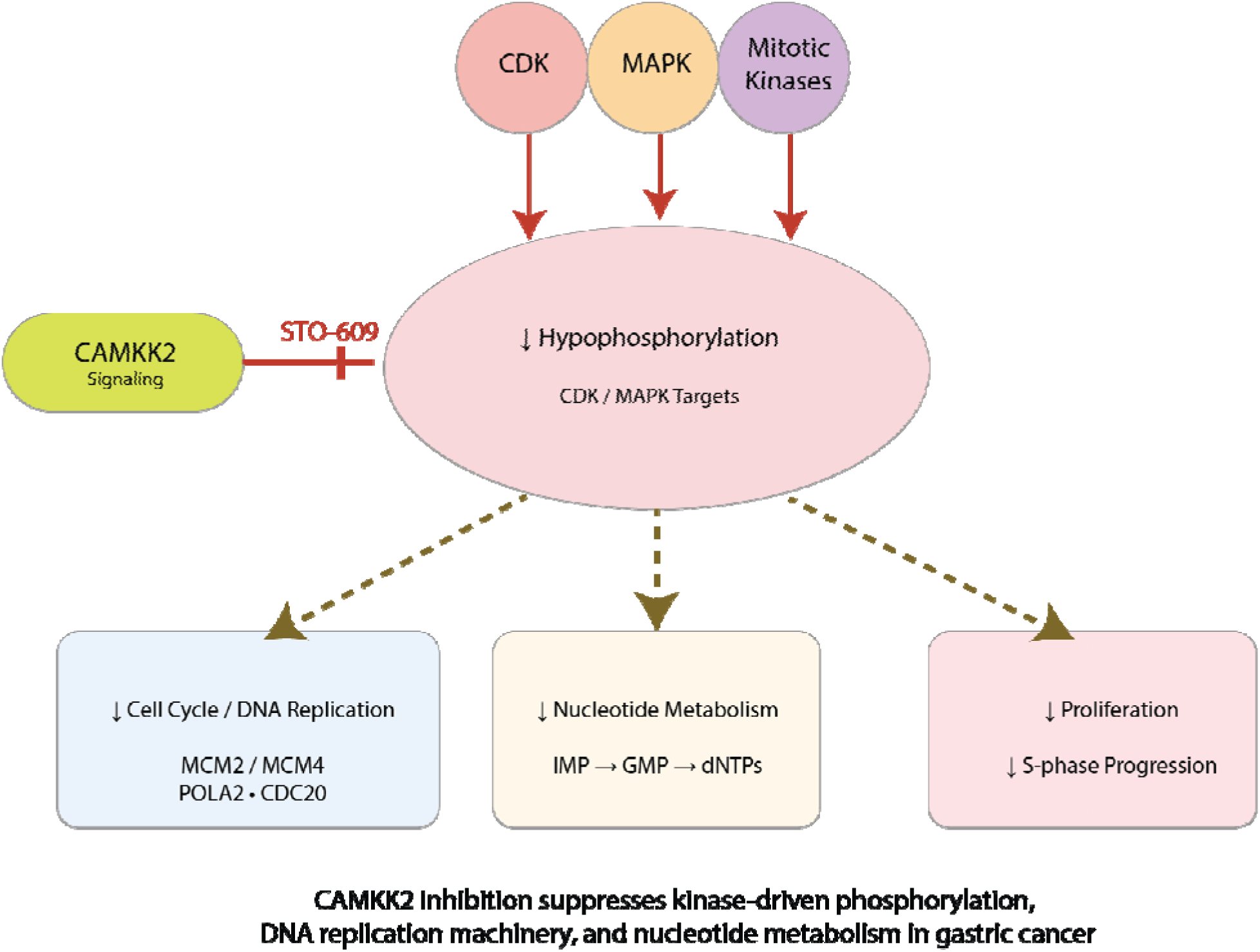
Proposed model of CAMKK2-mediated regulation of cell cycle and nucleotide metabolism in gastric cancer: Schematic representation of the effects of CAMKK2 inhibition in gastric cancer cells. Pharmacological inhibition of CAMKK2 using STO-609 leads to reduced phosphorylation of CDK, MAPK, and mitotic kinase targets, resulting in global hypophosphorylation of proteins involved in proliferative signaling. This suppression is associated with downregulation of cell cycle and DNA replication machinery (e.g., MCM2, MCM4, POLA2, CDC20), decreased nucleotide metabolism (IMP → GMP → dNTPs), and impaired S-phase progression. Collectively, these changes lead to reduced cellular proliferation and induction of replication stress.

In parallel, metabolomic profiling demonstrated significant depletion of nucleotide intermediates, including purine and pyrimidine metabolites required for DNA synthesis. Integration of metabolite and protein datasets further established a functional link between impaired nucleotide metabolism and reduced activity of replication machinery. These molecular alterations collectively culminated in a pronounced anti-proliferative phenotype, characterized by reduced cell number, impaired S-phase progression, and accumulation of multinucleated cells, indicative of mitotic failure. This phenotype is consistent with replication stress arising from insufficient nucleotide availability and defective replication signaling.

Together, these findings support a model in which CAMKK2 functions as a central regulator of proliferative capacity by coordinating kinase signaling, nucleotide metabolism, and DNA replication machinery. Inhibition of CAMKK2 disrupts this regulatory axis, leading to replication stress, cell cycle arrest, and impaired cellular proliferation in gastric cancer cells (**Figure 7**).

## Discussion

The present study provides a comprehensive multi-omics framework that positions CAMKK2 a a central regulator of proliferative signaling and metabolic homeostasis in gastric cancer. While CAMKK2 has been previously implicated in cancer metabolism and growth regulation, particularly through AMPK-dependent and independent pathways [23, 61], its direct role in coordinating nucleotide metabolism with DNA replication and cell cycle progression has remained largely unexplored. Our findings extend this paradigm by demonstrating that CAMKK2 integrates kinase signaling and metabolic pathways to sustain DNA synthesis and cellular proliferation.

A key observation of this study is the widespread suppression of phosphorylation-dependent signaling following CAMKK2 inhibition. Phosphoproteomic analysis revealed reduced phosphorylation of proteins involved in DNA replication, chromatin organization, and mitotic progression, accompanied by decreased activity of CDK and MAPK-associated signaling pathways. These findings are consistent with previous studies demonstrating that CDKs and MAPKs are critical regulators of S-phase entry, replication origin firing, and mitotic progression [58, 62, 63]. Notably, CAMKK2 has been reported to modulate kinase signaling networks through upstream regulation of Ca2+-dependent pathways [19], suggesting that its inhibition may disrupt multiple kinase cascades simultaneously. Our data therefore position CAMKK2 as an upstream modulator of kinase-driven proliferative signaling.

In parallel with signaling alterations, we observed pronounced metabolic reprogramming characterized by depletion of nucleotide intermediates. Cancer cells are known to rely heavily on de novo nucleotide biosynthesis to support rapid proliferation [34, 35, 37]. Disruption of nucleotide metabolism has been shown to induce replication stress due to insufficient dNTP pools required for DNA synthesis [64–66]. In agreement with these reports, our metabolomic analysis revealed significant downregulation of purine and pyrimidine metabolites, including intermediates essential for DNA replication. Importantly, proteomic data demonstrated concurrent downregulation of enzymes such as DUT, GMPR, and DNA polymerases, further reinforcing the link between metabolic insufficiency and impaired replication machinery.

The convergence of impaired kinase signaling and nucleotide depletion provides a mechanistic explanation for the replication stress phenotype observed upon CAMKK2 inhibition. Replication stress is a hallmark of cancer cells and can arise from oncogene activation, nucleotide depletion, or defects in replication machinery [40, 67, 68]. Our functional data demonstrating multinucleation and mitotic defects are consistent with previous studies showing that replication stress leads to aberrant mitosis, cytokinesis failure, and genomic instability [69, 70]. Thus, CAMKK2 inhibition appears to phenocopy classical replication stress–inducing conditions by simultaneously disrupting both signaling and metabolic inputs required for DNA synthesis.

Interestingly, enrichment analysis using Drug GeneSetDB revealed overlap between CAMKK2 inhibition signatures and those induced by nucleotide metabolism inhibitors such as hydroxyurea. Hydroxyurea is known to inhibit ribonucleotide reductase, leading to depletion of dNTP pools and replication arrest [69, 70]. The similarity between these signatures suggests that CAMKK2 inhibition may induce a comparable metabolic state, thereby reinforcing the concept that CAMKK2 is essential for maintaining nucleotide homeostasis. This finding also highlights the potential of CAMKK2 targeting as a strategy to induce replication stress selectively in cancer cells.

Our integrative protein–metabolite and docking analyses further provide structural and functional insights into the coupling between metabolic intermediates and enzymatic activity. Favorable binding interactions between nucleotide metabolites and key enzymes such as DUT and GMPR support the notion that metabolite availability directly influences enzyme function. Emerging evidence suggests that metabolic intermediates can regulate enzyme activity and stability through substrate availability and feedback mechanisms [71, 72]. Disruption of these interactions upon CAMKK2 inhibition may therefore contribute to the observed defects in nucleotide metabolism and DNA replication. From a broader perspective, our findings align with the growing concept that cancer cell proliferation is governed by tight coordination between signaling pathways and metabolic networks [73–75]. CAMKK2 appears to function as a critical node within this regulatory network, linking calcium signaling to metabolic and proliferative outputs. Given that CAMKK2 has also been implicated in other cancer types, including prostate and liver cancer [76, 77], the mechanisms described here may represent a more generalizable feature of CAMKK2-driven oncogenic processes.

Despite the strengths of this study, including multi-omics integration and functional validation, certain limitations should be acknowledged. While our data strongly support a role for CAMKK2 in regulating replication and metabolism, the direct substrates and downstream effectors of CAMKK2 remain to be fully elucidated. Additionally, in vivo validation and clinical correlation will be important to establish the translational relevance of these findings.

In conclusion, our study identifies CAMKK2 as a central integrator of kinase signaling, nucleotide metabolism, and DNA replication in gastric cancer. Inhibition of CAMKK2 disrupts this regulatory axis, leading to replication stress, mitotic defects, and impaired proliferation. These findings provide new mechanistic insights into CAMKK2 function and highlight its potential as a therapeutic target for inducing metabolic and replication vulnerabilities in cancer cells.

## Abbreviations

CAMKK2: Calcium/calmodulin-dependent protein kinase kinase
GC: Gastric cancer
LC-MS/MS: Liquid chromatographytandem mass spectrometry
GO: Gene Ontology
KSEA: Kinasesubstrate enrichment analysis
MAPK: Mitogenactivated protein kinase
CDK: Cyclin-dependent kinase
ERK: Extracellular signal-regulated kinase
AKT: Protein kinase B
ATM: Ataxia telangiectasia mutated
ATR: Ataxia telangiectasia and Rad3-related
PRKDC: Protein kinase DNA-activated catalytic subunit
PCA: Principal component analysis
FDR: False discovery rate
PTM: Post-translational modification
Fe-NTA: Ferric nitrilotriacetic acid
ACN: Acetonitrile
TFA: Trifluoroacetic acid
TEABC: Triethylammonium bicarbonate
SEM: Standard error of the mean

## Ethics approval and consent to participate

Not applicable

## Data Availability Statement

Data related to the study are provided in the manuscript and supplementary rest of the data and can be provided request.

## Author Contributions

P. K. M. conceived the idea, designed the experiments and interpretation of data, and critically reviewed and edited the manuscript. M.A.N conceived the idea, designed the experiments along with P.K.M. performed experiments and data analysis, drafted the manuscript, and prepared figures. N.C. Performed metabolomics data analysis drafted figures. N.D. Performed docking data analysis.

## Acknowledgements

The authors gratefully acknowledge Yenepoya (Deemed to be University) for providing the infrastructure and state-of-the-art mass spectrometry facility required to conduct this study exclusively within our institution. We also acknowledge the support provided by the Department of Biotechnology (DBT) through the National Facility grant under the project “Skill Development in Mass Spectrometry-based Metabolomics Technology BIC” (BT/PR40202/BTIS/137/53/2023).

